# Remembering what is relevant: how is goal-directed memory reactivation supported by attentional selection?

**DOI:** 10.1101/2023.03.15.532752

**Authors:** Melinda Sabo, Edmund Wascher, Daniel Schneider

**Affiliations:** Leibniz Research Centre for Working Environment and Human Factors, Ardeystraße 67, 44139 Dortmund, Germany

**Keywords:** episodic long-term memory, selective memory retrieval, attention, EEG, multivariate pattern analysis

## Abstract

Goal-directed memory reactivation involves retrieving the most relevant information for the current behavioral goal. Previous research has linked this process to activations in the fronto-parietal network, but the underlying selection mechanisms remain poorly understood. The current electroencephalogram (EEG) study investigates attention as a possible mechanism supporting goal-directed retrieval. Participants learned associations between objects and two screen locations in an episodic long-term memory experiment. In a following phase, we changed the relevance of some locations to simulate goal-directed retrieval. This was subsequently contrasted to a control condition, in which the original associations remained unchanged. Behavior performance measured during final retrieval revealed faster and more confident responses in the selective vs. neutral condition. At the EEG level, we found significant differences in decoding accuracy, with above-chance effects in the selective cue condition but not in the neutral cue condition. Additionally, we observed a stronger posterior contralateral negativity and lateralized alpha power in the selective cue condition. Overall, these results suggest that attentional selection enhances task-relevant information accessibility, emphasizing its role in goal-directed memory retrieval.

## 1. Introduction

Our episodic long-term memory (eLTM) has an impressive capacity to encode numerous details associated with a particular event. However, during retrieval, only a subset of the originally encoded information may be relevant to the current goal. The phenomenon of retrieving only the task-relevant information is known as goal-directed memory reactivation^1^. At the neural level, goal-directed memory reactivation was associated with activations of the fronto-parietal network, which were argued to represent retrieved memories in accordance with behavioral goals^1,2^. Nevertheless, the mechanisms underlying the selection of information in the service of goal-directed memory retrieval are still not well understood.

One important process that is likely to play a role in this context is attention. Theoretical models, such as the Attention to memory (AtoM) account posits that the role of top-down attention during long-term memory retrieval is to keep the current goal active, whereas the function of bottom-up attention is to signal when a change in attentional focus is required^3,4^. Furthermore, a recent study found that brain activity patterns recorded during an attention manipulation task more closely resembled the retrieval state than the encoding state of a long-term memory task. The author concluded that this highlights the presence of attentional states during long-term memory retrieval^5^. In a different study, Logan and colleagues^6^ reached comparable conclusions. Based on their behavioral findings, they suggested that long-term memory retrieval can be understood in terms of internal attentional processes. Finally, in a previous investigation, we demonstrated that the retrieval of relevant vs. irrelevant features from long-term memory requires different attentional processes (i.e., attentional selection and inhibition), which are reflected in distinct lateralized alpha-beta power modulations^7^.

Based on these previous findings, an important question that remains unanswered is whether and how attentional processes support the selection of task-relevant information in the service of goal-directed retrieval. The current electroencephalogram (EEG) study aims to address this issue. Answering this question has significant implications for the theoretical differentiation between goal-directed and incidental memory reactivation. The letter process refers to the reactivation of all features originally encoded as part of an episode^1^. Previous studies have suggested a distinction at the neural level, with the former being supported by frontal and parietal areas, and the latter by the medial temporal cortex^1,8^. The current study contributes to this debate by exploring whether the involvement of attentional selection is another feature that distinguishes the two forms of long-term memory retrieval.

To answer our main question, we designed an experiment in which we introduced a condition requiring goal-directed memory retrieval. This was subsequently contrasted with a control condition, in which all originally encoded information had to be retrieved. The task consisted of three phases: encoding, cueing, and retrieval phase. During the encoding phase, we asked participants to learn associations between an object and two spatial locations. Next, during the cueing phase, we changed the relevance of some of the initially encoded locations. Thus, for certain objects (selective cue condition), a cue indicated that only one of the two locations is relevant. Consequently, the selective cue condition is considered to be a proxy of goal-directed memory retrieval. In contrast, the control condition featured a so-called neutral cue, which indicated that both locations initially associated with the object were still relevant. During the final retrieval phase, participants had to report the task-relevant location associated with each object (see Figure 1).

**Figure 1.**
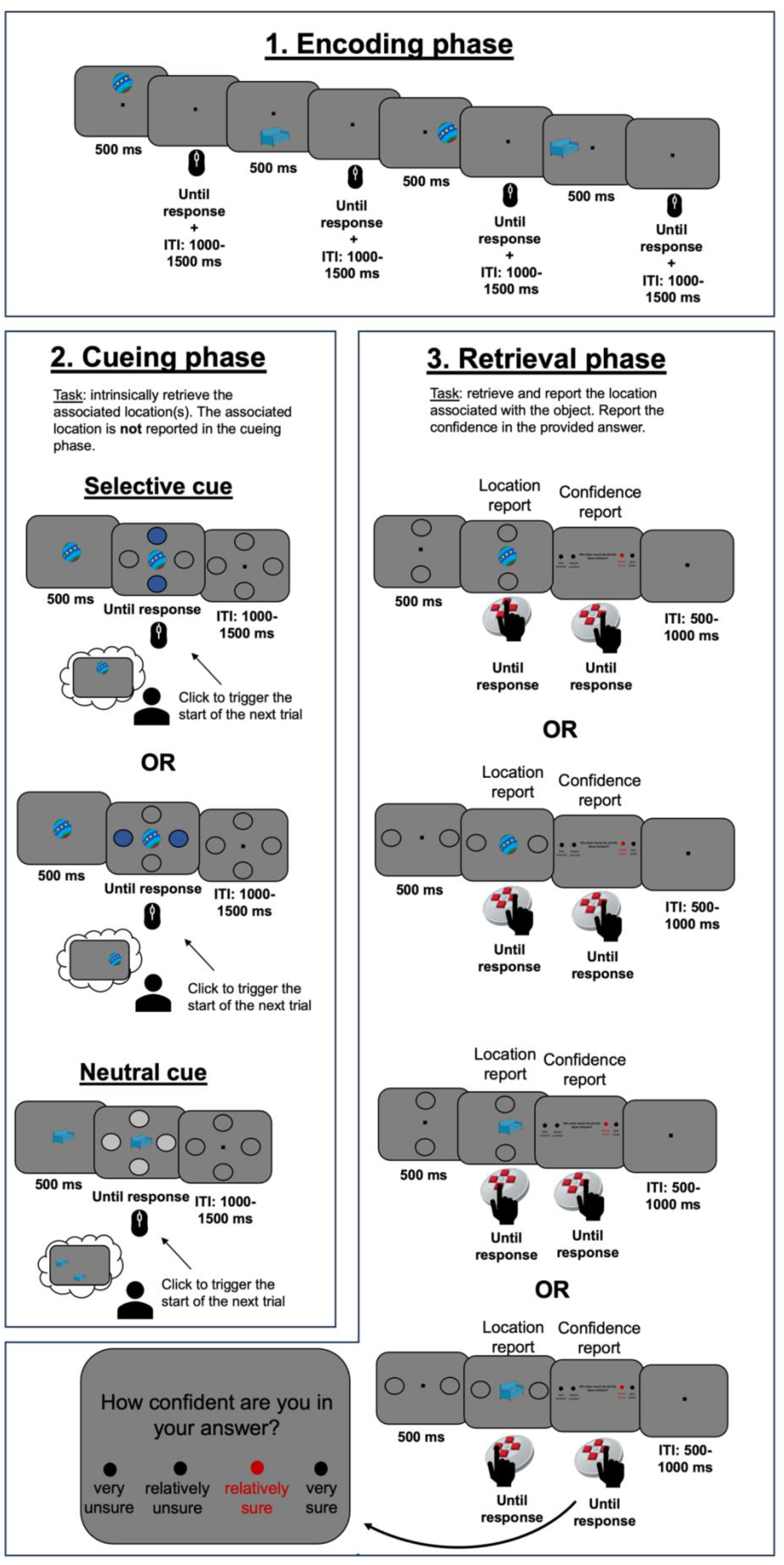
Schematic illustration of the experimental design. In the encoding phase, participants learnt associations between everyday objects and two locations on the screen. Each trial was self-paced and after participants finalized the encoding of the relevant association, they had to confirm it via a mouse click, which led to the start of the next trial. Between trials, we introduced an inter-trial interval (ITI) randomly varied between 1000-1500 ms. In the second phase (i.e., cueing phase), the objects from the previous phase were again presented along with a selective or neutral cue. Here, participants’ task was to either selectively think about one of the two locations associated with the object or to think about both originally encoded locations. Similar to the first phase, responses (via a computer mouse) were only necessary to confirm that participants completed the process of selectively thinking about the task relevant association. No locations had to be reported during the second phase of the task. Between trials, we introduced an ITI randomly varied between 1000-1500 ms. Finally, in the retrieval phase, participants were centrally presented with the objects from the previous phases and had to report the associated location. The two circles presented before the object onset indicated the to-be-reported positions: the one situated along the vertical or the horizontal axis. After the location report, participants were required to rate their confidence in the response, on a scale from one to four (1 = ‘Very unsure’; 4 = ‘Very sure’). The trial ended with an ITI randomly varied between 500-1000 ms. During the retrieval phase, a response device specifically designed for the current study was used.

As our main analysis procedure, we adopted a multivariate pattern analysis approach, in which we aimed to decode the locations associated with each object. Accordingly, if the EEG signal represents these location identities, this should result in above chance decoding performance^9^. Previous research has shown that locations held in working memory can be successfully decoded based on alpha power activity^10^. Furthermore, there is also evidence that information held in the focus of attention is decodable from the EEG patterns, while information outside the focus of attention is associated with below-chance decoding performance^11–13^. Based on these findings, we expect to find a difference in decoding accuracy between the selective and neutral cue conditions. More specifically, we predict higher decoding accuracy of the target location following the selective cue compared to the neutral cue.

Although previous research has shown that attended information yields above-chance decoding accuracy^11–13^, it is possible that condition differences obtained via decoding are not related to attentional processes. Therefore, the hypothesized effect needs to be confirmed by EEG correlates that are clearly related to attentional selection, such as lateralized alpha power (∼8-13 Hz)^14–17^ or the posterior contralateral negativity (PCN)^18–20^. Research has shown that directing attention to a lateral object or its memory representation results in a decrease in alpha power that is more pronounced in the hemisphere contralateral vs. ipsilateral to the location of the object^14–17,21^. Similarly, it has been argued that the PCN, an event-related potential (ERP) component, emerges as a stronger negativity contralateral vs. ipsilateral to the location of a lateral target stimulus^18–20^. Since in the context of the current long-term memory experiment we do not expect to obtain an asymmetry in the N2 time window, we will label our component as PCN. This pattern of activity is often referred to as the N2 posterior contralateral component (N2pc)^22^. In visual search tasks, these modulations were suggested to reflect target selection among distractors^18,19^, whereas in working memory the selection and prioritization of relevant items^20,21,23^. Based on these studies, we expect to find a difference in lateralized alpha power and PCN activity between the selective and neutral cue conditions. In line with the expected decoding results, we predict a stronger modulation of these EEG correlates in the selective compared to the neutral cue condition.

## 2. Results

### 2.1. Behavioral results

In order to investigate whether the use of the selective cue resulted in any behavioral benefit during the retrieval phase, we compared response accuracies (measured as the % of correct responses), the response confidence rating, and response times between selective and neutral trials. Our results revealed no significant differences in accuracy between the location report linked to the selective trials (*M_selective_* = 84.12 %; *SD_selective_* = 7.88 %) and the location report in the neutral trials (*M_neutral_* = 84.83 %; *SD_neutral_* = 8.55 %): *t*(29) = -0.61, *p* = 0.54, *d_av_* = 0.08, 95% CI [-3.04 1.62] (Figure 2a). When it comes to response times of all trials (irrespective of correctness), results showed that participants were significantly faster in reporting the location of an object occurring in the selective cue condition (*M_selective_* = 1425.99 ms; *SD_selective_* = 683.82 ms), as compared to the neutral cue condition (*M_neutral_* = 1560.89 ms; *SD_neutral_* = 685.51 ms): *t*(29) = -2.69, *p* = .01, *d_av_* = 0.19, 95% CI [-237.30 -32.49] (Figure 2b). When only the response times (RTs) of correct responses were considered, results suggested again that participants were faster in the selective, as compared to the neutral cue condition: *t*(29) = -3.13, *p* = .003, *d_av_* = 0.23, 95% CI [-255.90 -54.02] (*M_selective_* = 1336.32 ms; *SD_selective_* = 648.61 ms; *M_neutral_* = 1491.29 ms; *SD_neutral_* = 668.35 ms) (Figure 2c). The confidence ratings followed a similar pattern.

**Figure 2.**
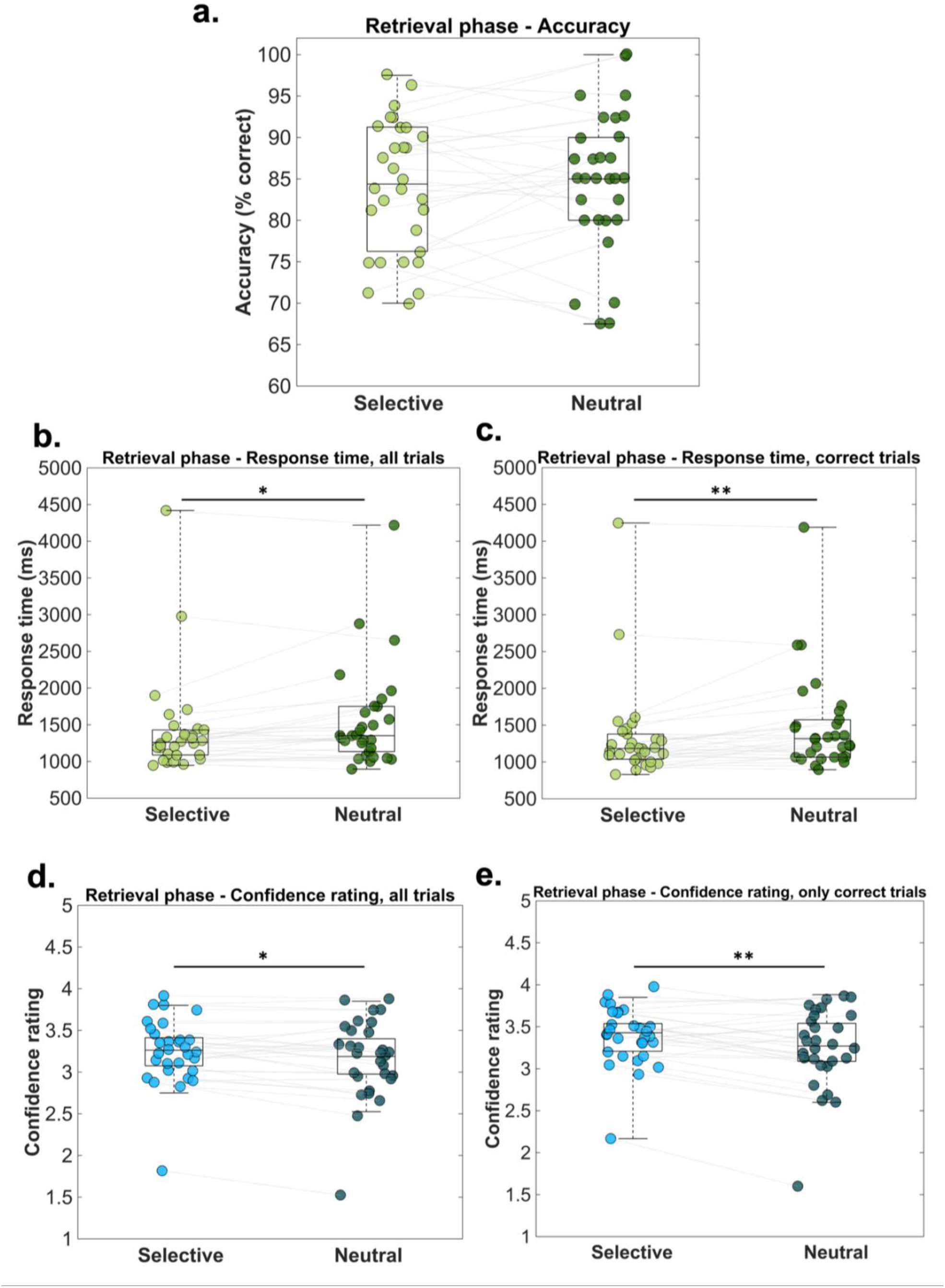
Behavioral results – retrieval phase. The scatterplots depict different behavioral parameters measured during the retrieval phase: a. average accuracy, b. average RT for all trials, c. average RT for only correct trials d. average confidence rating for all trials, and e. average confidence rating for only correct trials. The central mark on each boxplot indicates the median, and the bottom and top edges of the box mark the 25th and 75th percentiles. The whiskers are not extended to averages, which are considered outliers. Data points belonging to the same participant are connected via the grey lines. Statistical significance is marked with corresponding symbols: * *p* < 0.05; ** *p* < 0.01.

Accordingly, participants were more confident in their responses in the selective cue condition (*M_selective_* = 3.23; *SD_selective_* = 0.36) than in the neutral condition (*M_neutral_* = 3.15; *SD_neutral_* = 0.47): *t*(29) = 2.26, *p* = .03, *d_av_* = 0.19, 95% CI [0.00 0.15] (see Figure 2d). The same pattern was obtained when only correct trials were considered for this comparison (*M_selective_* = 3.37; *SD_selective_* = 0.33; *M_neutral_* = 3.24; *SD_neutral_* = 0.46): *t*(29) = 3.27, *p* = .002, *d_av_* = 0.32, 95% CI [0.04 0.21] (see Figure 2e). Overall, these results suggest a performance benefit associated with the selective cue in terms of response time and response confidence, but not in terms of accuracy.

### 2.2. Decoding analysis

Figure 3 illustrates the average decoding accuracy of the selective and neutral conditions. The classifier was trained to differentiate between target locations based on the scalp distribution of the EEG oscillatory power in the alpha frequency range (8-13 Hz; see further details in Methods). As a first step, we compared the decoding accuracy between the two conditions (time window of analysis: 500-2000 ms). This resulted in a significant cluster in the 1180-1360 ms time window (see gray area in Figure 3). Next, we contrasted the decoding accuracy to chance level. Our statistical procedure revealed a significant cluster ranging between 1140-1460 ms for the selective cue condition, but no statistically reliable effects for the neutral cue condition. Overall, we argue that these results support our hypothesis, which predicted a differential processing pattern between trials with a selective vs. a neutral cue.

**Figure 3.**
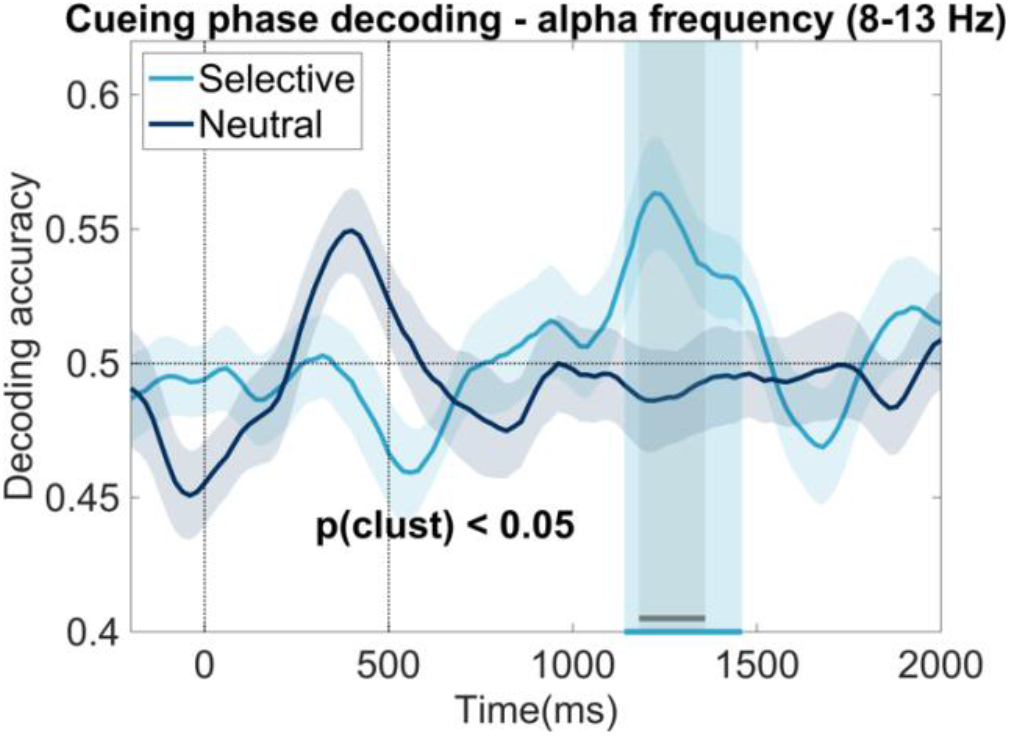
Decoding analysis (8-13 Hz activity) results – cueing phase. The figure shows the average decoding accuracy over time separately for neutral and selective conditions. The light blue shaded area shows the significant cluster obtained when decoding accuracy in the selective cue condition is contrasted to the chance level (significant cluster: 1140-1460 ms). The grey shaded area reflects the significant cluster obtained when contrasting the decoding accuracy of the two conditions (1180-1360 ms). The shaded area around the decoding accuracy time series depicts the standard error of the mean.

### 2.3. Posterior lateralized alpha power

Since our decoding results supported a differential decoding pattern between the selective and neutral conditions, we aimed to further investigate whether EEG correlates previously associated with attentional mechanisms are differentially modulated in the two conditions.

First, we explored lateralized oscillatory power in the alpha (8-13 Hz) frequency-band, which has been previously associated with attentional modulations in the perceptual and working memory domain ^15,16,24^. When it comes to our statistical comparison, we compared first the contralateral-minus-ipsilateral activity between the two conditions. Since the attentional modulation was expected to occur after the cue presentation (at 500 ms) and it was not anticipated to last longer than 1000 ms, the time window of analysis was restricted to 500-1500 ms. As indicated in Figure 4, our cluster-based permutation analyses did not reveal any significant results. Next, we contrasted the contralateral and ipsilateral power within each condition. Results for the selective cue condition revealed a significant cluster between 673-891 ms, while in case of the neutral cue condition, no significant differences were found (see Figure 4). Overall, these results suggest the presence of attentional selection processes during the selective cue condition.

**Figure 4.**
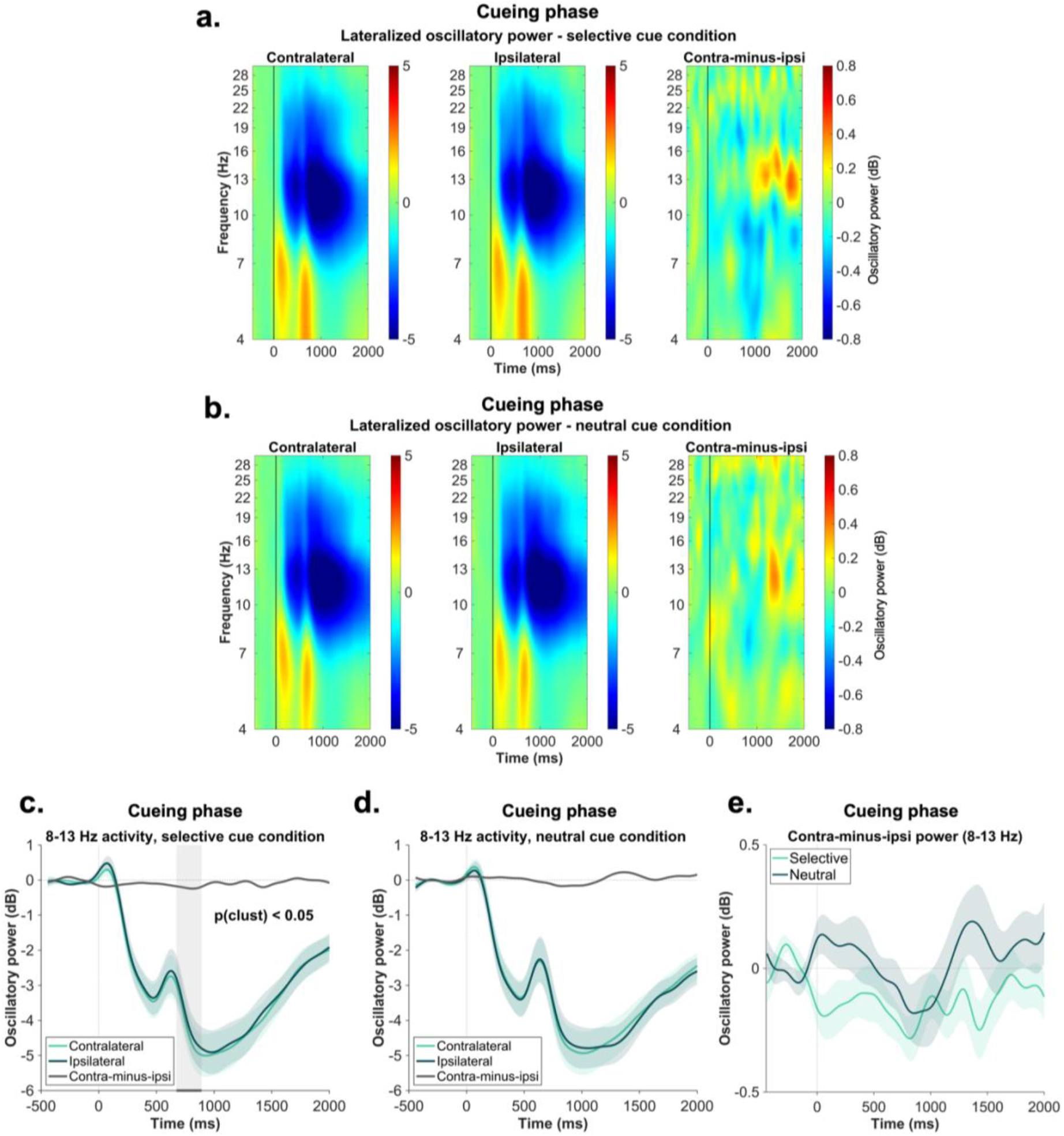
Lateralized alpha-beta power analysis results – cueing phase. Panel a depicts the time-frequency representation of the activity contralateral, ipsilateral and its difference relative to the lateral target positions in the selective cue condition (panel b for the neutral cue condition). The vertical line at 0 ms indicates the object onset. Panel c shows the corresponding line plots obtained by averaging the contralateral and ipsilateral activity across the 8-13 Hz frequency-band. The gray shaded area reflects the significant cluster obtained as a result of the cluster based permutation statistics: 673-891 ms. Panel d illustrates the alpha lateralization relative to the lateral locations in the neutral cue condition. Panel e depicts the comparison between the contralateral-minus-ipsilateral activity averaged across the alpha frequency range (8-13 Hz). The green shaded area around the individual difference waves represents in all figures the standard error of the mean.

### 2.4. Posterior ERP asymmetry

Beyond the alpha power modulations, we also investigated the PCN component, as a correlate of attentional selection (Figure 5a, 5b, 5c). First, we compared the contralateral-minus-ipsilateral activity between the two conditions. Similar to the lateralized alpha analysis, the time window was restricted to 500-1500 ms. Results indicated a significant cluster in the time window 966-1035 ms (see Figure 5b). Second, when the contralateral and ipsilateral activity was contrasted within the selective cue condition, our analysis revealed a significant cluster between 886-978 ms (see Figure 5a). When it comes to the comparison for the neutral cue condition, we did not find any significant differences. Overall, these results are in line with the decoding and alpha lateralization patterns and confirm that attentional processes contributed to the selective retrieval of information in the selective cue condition.

**Figure 5.**
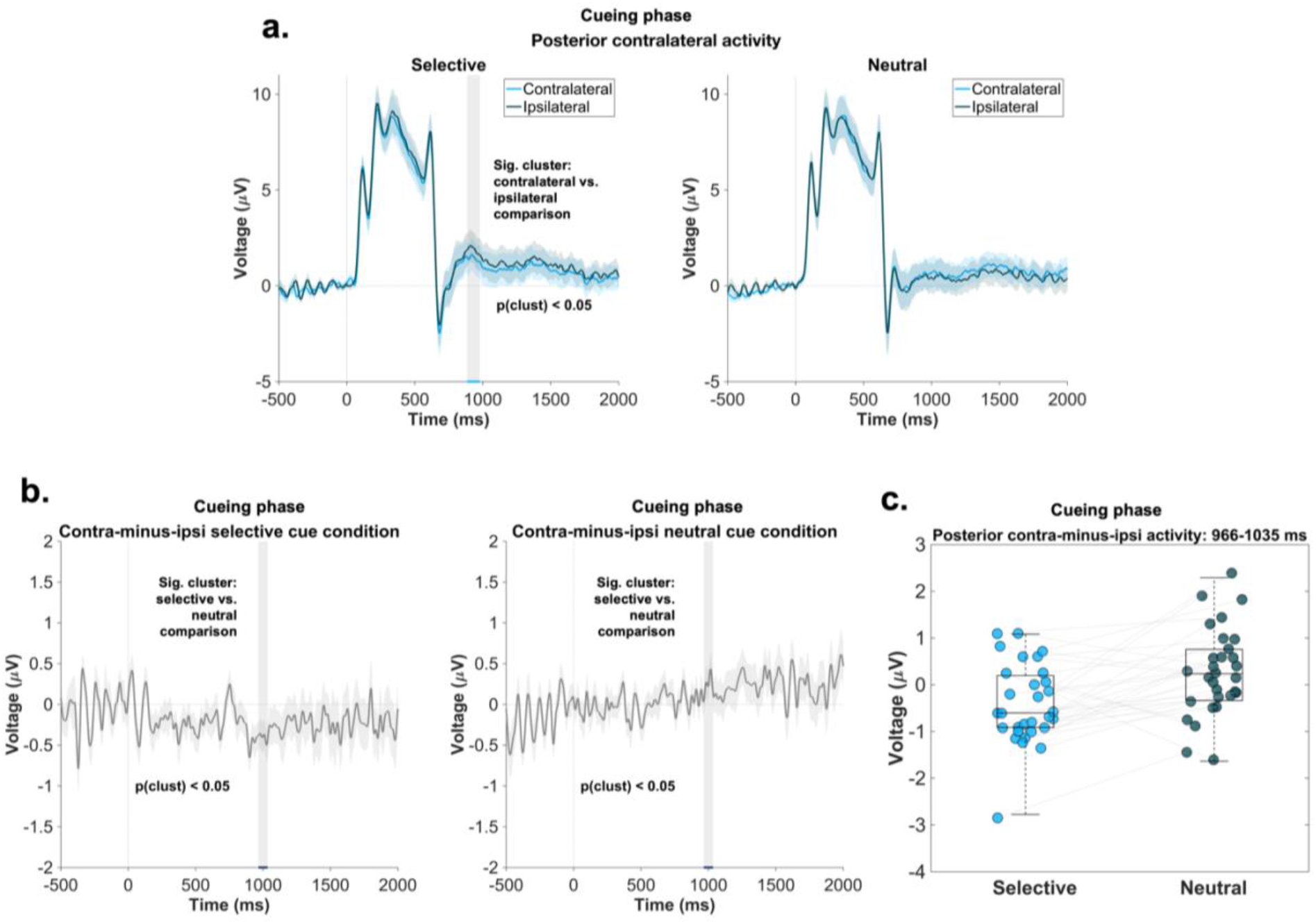
PCN results – cueing phase. Panel a. represents the posterior contralateral and ipsilateral activity relative to the lateral locations in the selective cue condition (left) and lateral locations in the neutral condition (right). The grey shaded area represents the significant time window obtained as a result of the statistical analysis conducted for the selective cue condition: 886-978 ms. The shaded blue areas depict the standard error of the mean. Panel b (left and middle) represent the contralateral-minus-ipsilateral activity in the selective and neutral cue condition. The shaded gray area illustrates the significant cluster obtained when the two conditions were contrasted: 966-1035 ms. The shaded gray area around the difference wave depicts the standard error of the mean. The right panel illustrates the scatterplot of the posterior contralateral-minus-ipsilateral activity averaged over the significant time window 966-1035 ms. The central mark on each boxplot indicates the median, and the bottom and top edges of the box mark the 25th and 75th percentiles. Data points belonging to the same participant are connected via the grey lines.

## 3. Discussion

In the current study, we explored attention as a mechanism for selecting information during goal-directed memory retrieval. We designed an eLTM experiment with a condition requiring goal-directed memory retrieval. After initial encoding, participants were cued on some trials to associate only one of two initially encoded locations with the objects (selective cue condition), serving as a proxy for goal-directed memory retrieval. We hypothesized that this condition would require attentional processes. In contrast, the neutral cue condition, where both locations remained relevant, did not require such selection and prioritization. We then compared brain activity recorded during these two conditions. We predicted condition differences in decoding accuracy (in line with^10–12)^ and in EEG correlates of attention, such as lateralized alpha power and the PCN component.

Behavioral results from the retrieval phase showed that participants responded faster and more confidently to objects from the selective cue condition (i.e., those paired with a selective cue in the second phase) compared to the neutral cue condition. However, we found no differences in accuracy between conditions. This discrepancy might be due to differences between the strength and accessibility of the representations. Specifically, selective cues may have increased the accessibility of stored information (indicated by faster reaction times and higher confidence) without affecting the strength of the representation (indicated by accuracy). Given our experimental design, it is unsurprising that we did not observe higher accuracy. After the initial encoding phase, participants received no feedback on the correctness of their internal retrieval during the second phase, leaving no opportunity to further reinforce learned associations.

The obtained behavioral patterns are consistent with some previous findings^25^, but they also contradict others^26–30^. First, our results align with a recent study that investigated the consequences of attentional selection during LTM retrieval^25^. This study suggested that informative cues, which direct participants’ attention to the location of the to-be-reported information, provided an advantage in response times compared to uninformative cues. Interestingly, this effect did not extend to accuracy. The authors proposed that the lack of accuracy difference might be due to ceiling effects in participants’ performance. This explanation might also partially apply to our results, as participants’ average performance during the final retrieval phase was relatively high (∼80%). Second, our findings appear to contradict previous studies on selective memory retrieval, which found memory enhancement for repeatedly retrieved items^26–30^. Importantly, those studies observed enhancement effects relative to a control condition where no retrieval occurred during the selective retrieval phase. In contrast, our control condition (i.e., neutral cue condition) required information retrieval but did not involve attentional selection. We chose this control condition to isolate potential attentional effects between conditions that were beyond mere retrieval effects. Consequently, since our design does not fully match the design of previous studies, we argue that the current results do not contradict these findings.

At the EEG level, our decoding analysis performed on alpha power revealed differences between conditions in the time window 1180-1360 ms after object presentation. Furthermore, we found that decoding accuracy was significantly above chance for the selective, but not for the neutral cue condition. This suggests that after the cue presentation, attention in the selective cue condition is more strongly focused on those locations where the association needs to be updated. Our results add to existing studies showing that, based on alpha power, inverted encoding models are able to track spatial locations retrieved from long-term memory^31^. Moreover, our interpretation is consistent with studies from the working memory field showing that information brought into the focus of attention is decodable, because it is maintained as a sustained pattern of neural activity^11,32,33^. In the current study, we provide evidence for a similar mechanism within the LTM domain. As such, items retrieved from LTM are similarly placed in an active state, where they undergo attentional processing, through which the relevant location information is selected, and the association is updated.

Our interpretation of attentional selection processes is further supported by the lateralized alpha and PCN results. We found a significant contralateral versus ipsilateral difference in the selective cue condition for both neural correlates. Additionally, a stronger PCN effect was observed in the selective versus neutral cue condition. As these EEG correlates have previously been shown to track attentional selection^14–21,34–36^ and the selective cue condition is regarded as a proxy for goal-directed memory retrieval, we argue that these results support the role of attentional selection processes in goal-directed memory retrieval.

The present findings have two important implications for the study of eLTM retrieval. First, the current study does not only bring evidence in favor of attentional selection supporting LTM retrieval in general^3–5,37^, but we specifically show that these processes play a particularly important role in goal-directed memory retrieval. Thus, based on our results, we suggest that attention is not a defining characteristic of all forms of retrieval, such as incidental memory reactivation, but it specifically defines goal-directed reactivation, enabling the selective access to information relevant for the current task demands. This adds to existing research efforts, which aim to dissociate incidental from goal-directed memory retrieval. For instance, Kuhl and colleagues^1^ indicated that goal-directed memory reactivation is supported by the fronto-parietal network, while incidental memory reactivation elicits medial temporal lobe activations. Similarly, Favila and colleagues^38^ indicated that the ventral parietal cortex represents retrieved features independent of their task-relevance, while the dorsal parietal cortex elicits activations only to relevant stimulus features.

Second, the current study has implications for on-going research exploring how the cognitive system handles partially overlapping memories, which are interfering with each other. This phenomenon is typically studied in paradigms in which participants learn two sets of overlapping associations (e.g., A-B, A-C). In a subsequent selective retrieval phase, they repeatedly retrieve only one out of the two pairs which becomes the target association (e.g., A-B), while the non-retrieved pair becomes the competing association. During the final retrieval phase, performance for all originally encoded associations is tested^26,27^. The typical observation in these paradigms is that the target memories are enhanced relative to a control condition, while performance for the competing memories decreases, which is referred to as retrieval-induced forgetting ^26–29,39–43^. At the neural level, it has been shown that competition is reflected in an increase in theta-band power^44–46^ and activations in the anterior cingulate cortex^47^ and the fronto-parietal network^48^. Moreover, a recent paper also suggests that a neural mechanism resolving the competition and interference is a distinct and separated coding of memories along the hippocampal theta rhythm^30^.

The experiment adopted in the current study shares some similarities with these paradigms, as we were interested in investigating the mechanisms underlying the repeated retrieval of the target information. However, the fate of the irrelevant item in the selective cue condition was beyond the scope of the current study. Overall, our results demonstrate that the target item’s accessibility increases as a result of attentional selection, thus providing compelling evidence that attentional processes are involved in interference resolution, and that they act at the level of the target representation. This interpretation is in accordance with the non-inhibitory account of interference resolution in retrieval-induced forgetting, which suggests that the selective retrieval of certain information has an effect on the target representation and as a consequence blocks the activation necessary for the competing, irrelevant information^39^. Since there are still ongoing debates about interference resolution during long-term memory retrieval, future research is needed to further elucidate how attention mediates these processes.

### 3.1. Conclusions

In summary, this study sought to elucidate whether and how attentional processes support the selection of task-relevant information in the service of goal-directed memory retrieval. Accordingly, we designed an experiment meant to capture relevance changes underlying goal-directed memory retrieval. Our decoding, oscillatory, and ERP results provided evidence for attentional selection in the selective cue condition as well as faster and more confident responses in the final retrieval phase. This indicates that attentional selection increased the accessibility of task-relevant information, highlighting its role in goal-directed memory retrieval. Two important implications have been discussed. First, the current results extend the distinction between incidental and goal-directed memory retrieval by suggesting that attentional selection only characterizes goal-directed reactivation. Second, in light of the current results, we propose attentional selection as a potential mechanism underlying interference resolution in long-term memory.

## 4. Methods

### 4.1. Participants

Initially 32 participants took part in the study. Since the instructions were provided in German and not all participants were native German speakers, two participants misunderstood the instructions. Instead of actively memorizing the associations, these two participants mistakenly believed that their task was to look briefly at each object. These two datasets were excluded from all analyses. Thus, the final sample contained data from 30 participants (age range: 18-29 years, *M* = 23.5 years, SD = 2.84 years) (16 females, 14 males). We based our sample size on previous studies that found reliable evidence for comparable attentional selection processes in working memory using EEG^17,24,49^. All participants were right-handed (as assessed by the Edinburgh Handedness Inventory^50^), had normal or corrected-to-normal vision and declared not suffering from any psychiatric or neurological disorder. Written informed consent was obtained from all participants before the start of the experiment. Participation was compensated with 10 Euros/hour or study completion credits (in case of psychology students). The study was conducted in accordance with the Declaration of Helsinki and was authorized by the ethics committee of the Leibniz Research Centre for Working Environment and Human Factors (Dortmund, Germany).

### 4.2. Procedure

Upon arrival, participants were provided with general information about the study. Subsequently, they filled in a demographic questionnaire, together with the Edinburgh Handedness Inventory ^50^. Afterwards, the EEG cap was prepared, and participants were seated in the dimly lit, sound-attenuated, and electrically shielded EEG chamber. The experiment was done on a 22-inch CRT monitor (100 Hz; 1024 × 768 pixels) and the viewing distance from the monitor was ∼145 cm. The task consisted of training followed by the three-phase eLTM experiment. Participants had a three-minute break between phases, with a three-minute distractor task in between. After completing the task, participants were given a follow-up questionnaire asking about the strategy used and any difficulties encountered. Half of the participants also took an intelligence test, but this is outside the scope of the current study. The duration of the whole procedure was ∼3-3.5 hours.

### 4.3. Experimental procedure

#### 4.3.1. Stimulus materials

For each participant, 120 objects were randomly chosen from a stimulus pool consisting of everyday objects^51^. The original stimulus pool contained 260 objects, but based on the image rating survey we conducted in a previous study^7^ we only considered a sub-sample of 240 objects, which had the best luminance, contrast, vividness, and were the most easily recognizable.

#### 4.3.2. Episodic-long-term memory task

The main task of the current experiment consisted of three main phases: encoding phase, cueing phase, and retrieval phase (see Figure 1). These phases were separated by a three-minute distractor task (counting backwards in steps of three from 500) and two breaks of three minutes each. This resulted in a total of nine minutes between phases one and two and between phases two and three. The experiment was preceded by a short training session to familiarize participants with the task. The training contained seven objects and was identical to the main experiment in terms of instructions.

In the encoding phase, participants were presented with an object (size: 4.3° × 2.9° or 2.9° × 4.3°) appearing on a gray background (RGB 128-128-128). The object was shown for 500 ms and could occur on four possible locations on the screen: top, bottom, left or right. Participants’ task was to memorize the association between the object and its presentation location, while fixating on the centrally presented black dot (size: 0.2° × 0.2°, distance between the object and the central fixation dot: 3°) (see Figure 1). Participants were instructed to use a specific strategy to remember the object-presentation location association. Specifically, after the object disappeared from the screen, they were instructed to visualize the object at the corresponding presentation location while maintaining their gaze on the central fixation dot. They were also instructed not to repeat the associations verbally. Memorization of an association was confirmed by clicking the mouse button with the index finger of the right hand, which led to the start of the next trial. If a participant took more than 15 s to memorize an association on several consecutive trials, the experiment-coordinating team verbally instructed the participant to proceed more quickly. During the encoding phase, each of the 120 objects was associated with two presentation locations: a lateral location (left or right) and a non-lateral location (top or bottom). To facilitate the learning process, each object-presentation location association occurred twice during encoding, resulting in a total of 480 trials (organized into eight blocks). Self-paced breaks were introduced between these blocks.

The cueing phase began with the central presentation of a previously seen object. After 500 ms, a set of four circles (size: 2°) appeared on the screen as a cue (distance between object and circles in visual angle: 3°). The reason for introducing a 500 ms time window between the object and the cue onset was to separate the memory reactivation from the attentional processes triggered by the cue presentation. The cues could be presented in one of the following configurations: (i) two blue circles (RGB: 50-85-150) along the vertical axis (i.e., top and bottom locations; selective cue); (ii) two blue circles (RGB: 50-85-150) along the horizontal axis (i.e., left and right locations; selective cue); (iii) four gray circles (RGB: 200-200-200) on top, bottom, left, and right locations (neutral cue; see also Figure 1). In the first case, participants’ task was to retrieve and selectively associate the object with the location along the vertical axis (selective cue condition). For example, if an object was associated with the top and left locations during encoding (see Figure 1), in the cueing phase participants had to associate only the top location with the respective object. In the second selective condition, only the location along the horizontal axis (i.e., left or right) was relevant. Finally, in the neutral condition, both locations originally associated with the object remained relevant. One third of the objects (40 in total) were assigned to the horizontal selective cue condition, another third (40 in total) to the vertical selective cue condition, and the remaining 40 objects to the neutral cue condition. Similar to the encoding phase, memorization of an association was confirmed by a right mouse click, which led to the start of the next trial. This mouse click was identical across trials, regardless of the condition or the object to be reported. During the second phase, participants were not required to report the associated location; their only task was to think about the cued location(s). In addition, participants who needed more than 15 s for several consecutive trials were verbally instructed to reduce the processing time. To facilitate learning, the 120 objects were presented three times together with the same cue (i.e., selective horizontal, selective vertical, or neutral). The cueing phase consisted of a total of 360 trials divided into six blocks.

Finally, the retrieval phase started with the presentation of two circles (size: 2°), together with a central fixation dot. These circles could appear either along the horizontal or the vertical dimension, in accordance with the cued dimension from the second phase. After 500 ms, one of the objects present in the previous phases appeared in the center, between the two circles (distance between the object and the circle in visual angle: 3°) (see Figure 1). Participants’ task was to report the location associated with the object, by clicking on the corresponding button of the response device with the index finger of the right hand. Since the task was generally demanding, we introduced a two-alternative forced-choice task. Accordingly, participants were either presented with the circles on top- and bottom- or on the left- and right locations. The dimension along which these circles appeared always corresponded to the cued axis from the cueing phase. In case of the neutral condition, the probed dimension (vertical or horizontal) was randomly selected, with the restriction that half of the trials cued the horizontal dimension, while the remaining half, the vertical one. The response device was equipped with four buttons, mapped onto the four possible locations on the screen (top, bottom, left, and right). After each response, participants were instructed to bring their index finger back to the middle location of the device. In addition, after the location report, participants were also required to rate their confidence in the provided response on a scale from 1 to 4 (1 = really unsure, 2 = relatively unsure, 3 = relatively sure, 4 = very sure).

### 4.4. Data analyses

All behavioral, EEG, and statistical analyses were conducted in MATLAB® (R2021b). As a first step, each participant’s dataset was divided according to the three phases of the task: encoding, cueing, and retrieval phase. The EEG analyses were conducted only on the cueing phase data.

#### 4.4.1. Behavioral analyses

In a first step, we assessed accuracy, response times, and confidence ratings during the retrieval phase, when participants had to report the location associated with each object. We separated the retrieval trials containing objects previously presented together with a selective cue from those containing objects presented together with a neutral cue during the cueing phase. In case of the confidence ratings and response times analysis, we first performed it for all trials regardless of correctness and then only for trials with correct responses^52,53^. Paired sample t-tests were used to examine the differences in each behavioral parameter between these two experimental conditions.

#### 4.4.2. EEG recording

The EEG data were collected with a passive 64 channel Ag/AgCl system (Easycap GmbH, Herrsching, Germany), with its electrode distribution following the extended 10/20 system ^54^. During recording, the signal was amplified by a NeuroOne Tesla AC-amplifier (Bittium Biosignals Ltd, Kuopio, Finland) and it was low-passed filtered at 250 Hz. The original sampling rate of the data was 1000 Hz and FCz acted as the reference electrode, while AFz served as the ground electrode. Throughout the session, the impedances were kept below 20 kΩ.

#### 4.4.3. Preprocessing

We preprocessed the data using the EEGLAB toolbox (version 14.1.2b)^55^ implemented in MATLAB®. In the following sections, we will first describe the preprocessing steps adopted in the case of lateralized alpha and ERP analysis (for an overview, see Figure 6), and then the specific details for the pipeline used for the decoding analysis (see the preprocessing output for each pipeline in Table 1).

**Figure 6.**
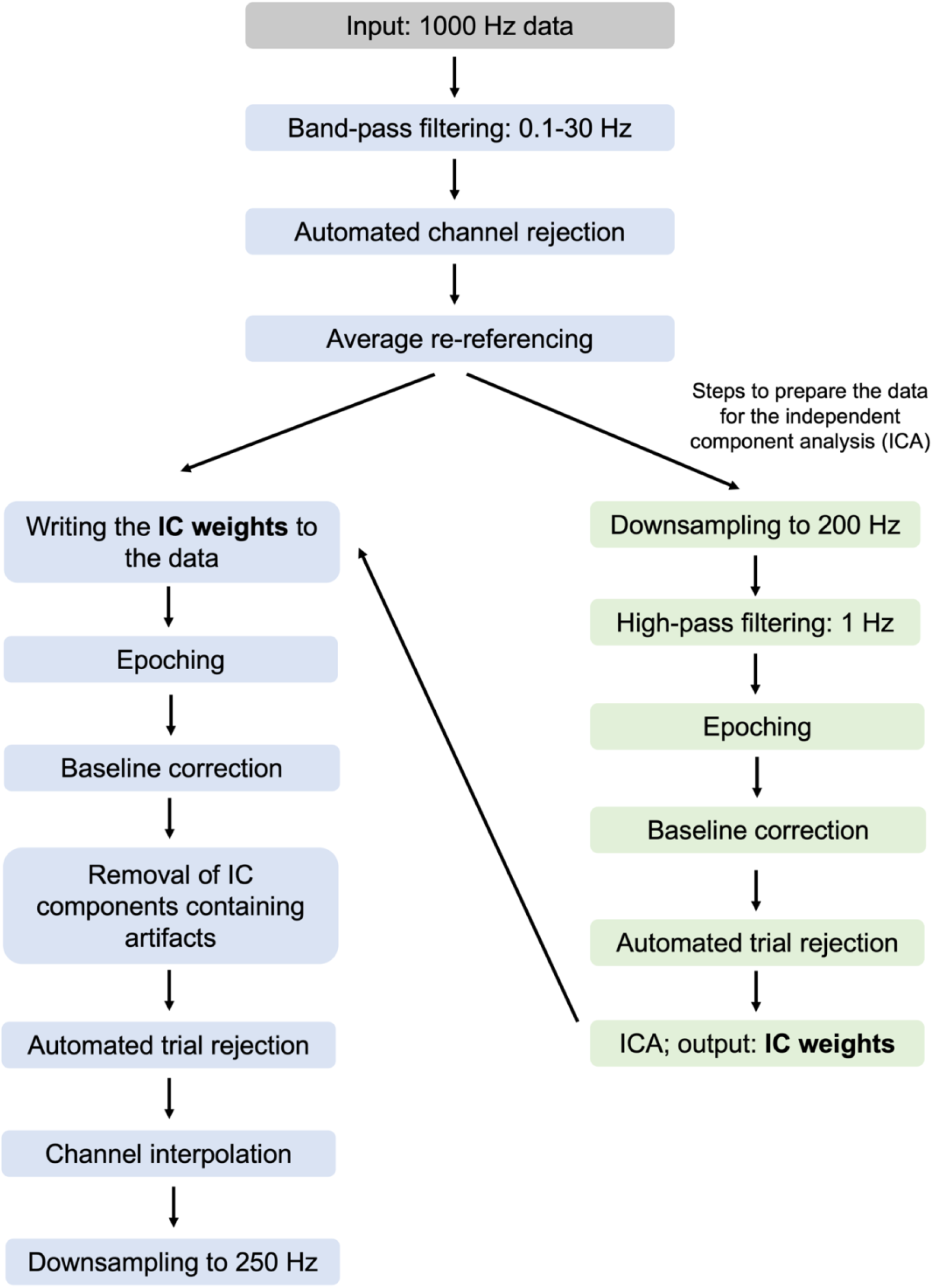
Schematic illustration of the preprocessing steps adopted for the lateralized alpha power and ERP analyses.

**Table 1.**
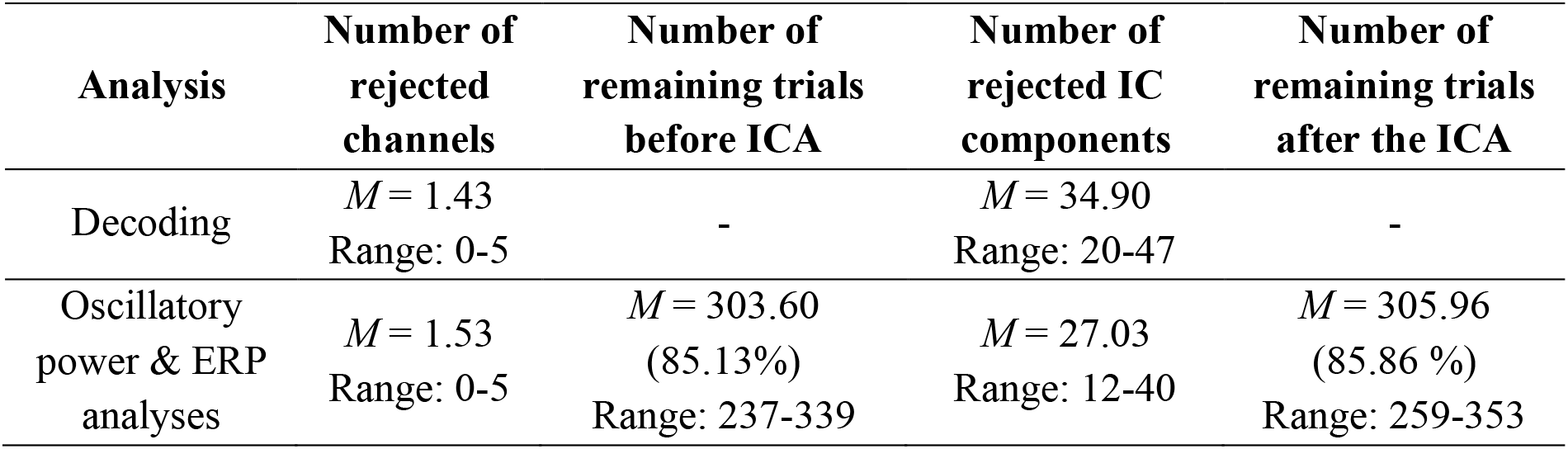
Overview of the outcome of each preprocessing pipeline.

As a first step in our preprocessing procedure, a 0.1 Hz Hamming windowed sinc FIR high-pass filter (filter order: 33001, transition bandwidth: 0.1 Hz, cutoff frequency at −6 dB: 0.05 Hz) and a 30 Hz low-pass filter (filter order: 441, transition bandwidth: 7.5 Hz, cutoff frequency at −6 dB: 33.75 Hz) were applied (*pop_eegfiltnew* function). Subsequently, channels containing high levels of noise were rejected using the automated channel rejection procedure adopted in EEGLAB (via the *pop_rejchan* function) and data were re-referenced to the average. The anterior channels capturing eye movements (Fp1/2, Fpz, AF3/4, AF7/8, Afz) were excluded from the channel rejection procedure in order to ensure an optimal detection of eye-movement-related components later in the independent component analysis (ICA). The following steps served to prepare the data for the ICA. Data were downsampled to 200 Hz and high-pass filtered using a 1 Hz Hamming windowed sinc FIR filter (*pop_eegfiltnew*, filter order: 661, transition bandwidth: 1 Hz, cutoff frequency at −6 dB: 0.5 Hz). Next, epochs ranging between -1000-3000 ms relative to the object onset were created, followed by baseline correction (time window of the baseline: -200-0 ms). Subsequently, trials containing extreme fluctuations were rejected via the automated trial rejection procedure of EEGLAB (i.e., *pop_autorej*; threshold: 500 μV, maximum % of rejected trials: 5%). Afterwards, the ICA was performed on the rank-reduced data (remaining number of channels minus 1 – obtained through the principial component analysis built into the *pop_runica* function). In order to identify the independent components (ICs) containing artifacts (e.g., eye movements, muscle activity, channel noise etc.), we used the ICLabel plug-in (version 1.3) ^56^. ICLabel operates with seven categories (brain, muscle, eye, heart, line noise, channel noise, and other noise) and in case of each IC, it assigns a probability for containing the type of noise described by these categories. For the current analysis, we rejected the ICs which were labelled with a 30% (or more) probability as an eye movement component, as well as those which were labelled with less than 30% probability as a brain component ^57^. Then, the weights obtained from the ICA were written back to the 1000 Hz data, which had been band-pass filtered and re-referenced to average. Next, epochs time-locked to the object onset (-1000-3000 ms) were created and data were baseline corrected (-200-0 ms baseline). Afterwards, the ICs marked for rejection were removed and trials still containing large fluctuations were rejected via the same automated procedure mentioned above (threshold: 1000 μV, maximum % of rejected trials: 5%). Finally, using the spherical spline method implemented in EEGLAB (i.e., *pop_interp*), missing channels were interpolated.

In the preprocessing of the data for the decoding analysis, no filtering was done (except for the 1 Hz low-pass filtering meant to prepare the data for the ICA), as it has been previously argued that such filtering can artificially inflate the decoding results^58^. Moreover, since decoding analyses are sensitive to number of trials in the input data, the trial rejection procedure was eliminated from this last pipeline. Since filtering and trial rejection were omitted, we decided to apply channel rejection to the full range of electrodes. Finally, we additionally downsampled the data by selecting every fifth time point.

#### 4.4.4. Time-frequency decomposition

After preprocessing, data were downsampled to 250 Hz and a complex Morlet wavelet convolution analysis was conducted. This procedure relies on convolving the data with a family of complex Morlet wavelets obtained by calculating the timepoint-wise dot product between complex sine waves of varying frequencies and Gaussians of different widths (as defined by the number of cycles). The complex Morlet wavelet family covered 26 logarithmically spaced frequencies between 4 and 30 Hz. The number of cycles started at four and increased in logarithmic steps until 11.25 (at the highest frequency). When it comes to the lateralized alpha analysis, we also adopted baseline normalization through decibel conversion. Consequently, the reported power values are in units of decibel (dB).

#### 4.4.5. Decoding procedure

The main aim of the present decoding procedure was to investigate whether different locations associated with the objects could be decoded based on the scalp distribution of the EEG power. In this context, we included EEG oscillatory power only from the alpha frequency-band (8-13 Hz)^59,60^. In our decoding procedure, we sought to test whether differences in decoding accuracy existed between the two conditions (i.e., selective cue and neutral cue condition). Accordingly, we focused on decoding the target location in these two conditions. More specifically, the classifier was trained to distinguish between: (i) left vs. right target locations (i.e., trials, in which the participant was supposed to selectively associate the object with the right location vs. trials, in which the participant was supposed to selectively associate the object with the left location); and (ii) top vs. bottom target locations. The reason for not including other contrasts, such as left vs. top or right vs. bottom was due to the selective cues differing in their perceptual characteristics. For example, in trials, in which the target location was left, two horizontal colored circles were shown, while in trials with a top target location, two vertical colored circles. Including such contrasts to our analysis would have led to decoding the identity of the cue (i.e., horizontal vs. vertical), instead of decoding the target location. To keep the analysis in the two conditions comparable, we included the same contrasts also for the neutral condition. Overall, the classification routine was ran two times for both conditions. We adopted the approach described by Bae and Luck^10^, who used a combined support vector machines and error-correcting output codes approach^61^, implemented into the *fitcecoc()* MATLAB® function. Testing and model predictions were done via the *predict()* function.

The input to our decoding procedure was constituted by the power values, resulting from the time-frequency decomposition. We included six frequency-bands between 8-13 Hz and 64 electrodes as features in our analysis. The input matrix of the classifier had a size of 384×4, where 384 results from the multiplication of the 64 channels by 6 frequencies and the second dimension reflects the number of trial averages used for training (see more details about trial averaging below). Classification was executed within each participant and timepoint separately. We focused on the EEG oscillatory activity between -200-2000 ms, containing 126 timepoints. As such, at each timepoint of an iteration, the trials belonging to each of the locations were divided randomly into three groups, from this point on referred to as blocks. An important aspect of this was to ensure that an equal number of trials associated with each location were assigned to each block. In cases where the number of trials could not be divided by three, these extra trials were removed. Within each block, data were averaged within each location label, thus resulting in three averages for each location. Here, averaging served the purpose of reducing noise, so classification was performed on trial averages rather than single trial data^10^. Averaging a random subset of trials containing different objects also rules out the possibility that the results are explained by decoding the object identity. The model training was done on 2/3 of these averages, while the test was conducted on the remaining 1/3. Since we performed a 3-fold cross-validation, each of these averages served once as a test dataset. Once this procedure was complete, trials were again shuffled and randomly re-assigned to one of the three blocks. Within these blocks, averages were again calculated and labeled as a training or a test dataset. This process was repeated in each of the 10 iterations. Once predicted labels were obtained for all timepoints and iterations within a participant, we calculated the decoding accuracy by comparing the predicted labels to the true ones. Decoding accuracy was subsequently averaged across iterations and in order to smooth these time series, a five-point moving time window was applied. Finally, decoding accuracies were averaged across participants. Given that we always contrasted two locations, the chance level was set to 50%.

When it comes to the statistical procedure, we first compared decoding accuracy between conditions, followed by a contrast relative to chance level, using the cluster-corrected sign-permutation test^11^. This procedure is based on generating a null distribution by flipping the sign of the values with a 50% probability, throughout 100000 iterations (function: *cluster_test_helper*). Subsequently, using this null distribution, significant clusters are determined in the actual data (function: *cluster_test)*. All significance thresholds were set to *p* < .05. Three different statistical comparisons were conducted: first, we contrasted the decoding accuracy between the two conditions, and then we compared the decoding accuracy obtained in each condition to the chance level. In case of the latter analysis, we performed one-tailed, cluster-corrected sign permutation tests because decoding accuracy was hypothesized to be higher than the chance level. All comparisons were restricted to the time window 500-2000 ms. The choice of the lower limit was based on the assumption that any potential attentional effect reflected in the differential activity will occur after the cue presentation (i.e., after 500 ms). The upper limit of the time window was selected based on the average response time, i.e., ∼2300 ms. The last 300 ms were omitted, because we did not want to capture response-related motor activity.

#### 4.4.6. Posterior lateralized alpha power analysis

Since our decoding results revealed differences between conditions, we also investigated EEG correlates of attention selection. Our first approach focused on lateralized alpha oscillatory power. Here, we calculated the average contralateral, ipsilateral, and contralateral-minus-ipsilateral activity relative to the lateral target location in the selective and neutral cue conditions. Importantly, we excluded trials, in which the selective cue marked the vertical dimension, and thus the target location was non-lateralized. This activity was then averaged across a posterior cluster, PO7/8, PO3/4, O1/2, chosen based on previous investigations^7,24^. In line with our hypotheses, we compared the contralateral-minus-ipsilateral activity between conditions and subsequently contrasted the asymmetry associated with each condition to zero. As a statistical procedure, we adopted a cluster-based permutation analysis, restricted to the time window 500-1500 ms. The main assumption in choosing this time window was that any attentional effect would occur after the cue presentation (at 500 ms) and would not last longer than 1000 ms^7^.

In the cluster-based permutation procedure, each voxel of the original time-frequency data was compared between conditions using paired-sample t-tests. This resulted in a matrix of *p*-values, containing 200 values (corresponding to 200 time points). If neighboring voxels containing *p*-values exceeding the significance level (i.e., 0.05) were found, the cluster’s size was saved. As a next step, the condition labels (i.e., contralateral or ipsilateral) were randomly exchanged within each dataset and paired sample t-tests were again conducted for each voxel of the time-frequency data. As before, clusters of significant *p*-values were identified and those containing the largest number of points were saved for the respective iteration. This procedure of re-shuffling the condition labels and identifying the largest cluster was repeated through 10000 iterations. This way, we could obtain a distribution of maximum cluster sizes, necessary to obtain the cutoff based on which significance is determined. Subsequently, if the cluster(s) identified in the original comparison were larger than the 95^th^ percentile of the cluster size distribution, they were considered to be significant.

#### 4.4.7. Posterior ERP analysis

In our ERP analysis, we aimed to further investigate the PCN component as a marker of attentional selection. As a first step in our analyses, we calculated the average contralateral, ipsilateral, and contralateral-minus-ipsilateral activity relative to the lateral target location in the selective and neutral conditions. Similar to the alpha lateralization analysis, we excluded trials, in which the selective cue marked the vertical dimension, and thus the target location was non-lateralized. The activity was obtained from PO7/8 electrodes^20^. Next, in order to perform a statistical comparison, we contrasted the contralateral-minus-ipsilateral activity between conditions using the same cluster-based permutation procedure described above (same time window ranging between 500-1500 ms). Finally, the contralateral and ipsilateral activity within each condition was also compared.

#### 4.4.8. Inferential statistics and effect sizes

As a measure of effect size, we used Cohen’s d_av_^62^. If not otherwise specified, the threshold for statistical significance was set to 0.05.

## Acknowledgments

The authors would like to thank Tobias Blanke for the offered technical support; Karin Lukaszewski for the help with participant recruiting; Pia Deltenre and Barbara Foschi for the lab organisation, and all the students, for contributing to the process of data collection: Joel Althoff, Lara Bleckmann, Nina Cialone, Quang Dang, Patrick Frenken, Kagan Kemerioglu, Vivek Mishra, Vivien Szczepanski, Stefan Weber.

## Author contribution

M.S.: data collection and data analysis. M.S. and D.S.: conceptualisation and writing the original draft. D.S.: project supervision and acquiring the necessary funding. E.W.: supervision and writing (review & editing).

## Open practices and data availability statement

The experiment was not preregistered. The raw data and all the scripts used for the reported analyses will be publicly available after publication on Open Science Framework (OSF): https://osf.io/p93gm/?view_only=44c1ea1c47d0436795e70dd83c89d752

